# Rapid reconstitution and kinase-controlled regulation of algal pyrenoid condensates in engineered *Nicotiana benthamiana*

**DOI:** 10.64898/2026.07.20.739623

**Authors:** Yana Kazachkova, Jason I. Punskovsky, Ashwani K. Rai, Eric Franklin, Martin C. Jonikas

## Abstract

The pyrenoid is an organelle found in nearly all algae that concentrates CO₂ around Rubisco to enhance photosynthetic carbon fixation. Engineering a functional pyrenoid into C₃ plants is a promising route to improve plant photosynthetic performance and yields. However, progress has been limited by the lack of a plant platform for rapidly testing candidate components. Here, we engineered *N. benthamiana* to make its chloroplast Rubisco holoenzyme compatible with *C. reinhardtii* pyrenoid proteins. Disruption of native *N. benthamiana RBCS* genes and complementation with the *C. reinhardtii RBCS2* generated a Rubisco-recombinant background where transient expression of the Rubisco linker protein EPYC1 was sufficient to drive the formation of Rubisco–EPYC1 condensates. Co-expression with the algal pyrenoid kinase KEY1 shifted the multi-condensate state toward a single condensate per chloroplast by altering EPYC1 phosphorylation *in planta*, recapitulating *C. reinhardtii* pyrenoid regulation. Together, these results establish engineered *N. benthamiana* as a tractable platform for rapidly characterizing candidate pyrenoid proteins and provide a step toward reconstructing pyrenoid architecture and regulation in plants.

## Introduction

Photosynthetic carbon assimilation in many crop plants is limited by the performance of the CO_2_-fixing enzyme Rubisco (Ort et al., 2015; Bouvier et al., 2024). In C₃ plants, Rubisco’s poor performance arises from low CO₂ availability at the site of carboxylation, resulting in decreased catalytic rate and frequent wasteful oxygenation reactions with Rubisco’s alternative substrate, O₂ (Tcherkez et al., 2006). To enhance Rubisco’s performance, some organisms have evolved CO₂-concentrating mechanisms (CCMs) that elevate CO₂ concentration around Rubisco (Raven et al., 2008; Sage et al., 2012). There is currently significant interest in understanding how these CCMs work and engineering them into plants to enhance crop performance (McGrath and Long, 2014; Hennacy and Jonikas, 2020; Fei et al., 2022). A functional CCM has the potential to enhance crop biomass, nitrogen use efficiency, water-use efficiency, and temperature resilience (Vogan and Sage, 2011; Long et al., 2015; Ainsworth and Long, 2021; Bernacchi et al., 2025).

The green algal pyrenoid-based CCM is particularly promising for engineering because of its unique characteristics. Unlike C₄ photosynthesis, which typically requires extensive modifications to leaf anatomy (Kajala et al., 2011), the algal CCM operates at the level of a single cell, simplifying the engineering required to reconstitute it in plants. Additionally, individual pyrenoid components are natively encoded in the nuclear genome, and most localize correctly when expressed in plants without modification; others have been shown to require only the addition of a known transit peptide for correct targeting (Atkinson et al., 2016). Among pyrenoid-based CCMs, the one from the green alga *Chlamydomonas reinhardtii* is the most promising target for engineering into plants, both because it is the best-understood and because *C. reinhardtii* is relatively phylogenetically close to land plants, increasing the likelihood that its components will be compatible with plant systems (Leliaert et al., 2012; Atkinson et al., 2016; He et al., 2023).

At its core, the *C. reinhardtii* pyrenoid contains a liquid-like biomolecular condensate, called the matrix (Freeman Rosenzweig et al., 2017), that houses densely packed Rubisco (Vladimirova et al., 1982; Lacoste-Royal and Gibbs, 1987). The matrix forms through phase separation driven by multivalent interactions between Rubisco and the intrinsically disordered linker protein EPYC1 (Mackinder et al., 2016; Wunder et al., 2018; He et al., 2020; Barrett et al., 2021). The matrix is traversed by specialized thylakoid membrane tubules that deliver concentrated CO₂ (Pronina and Semenenko, 1990; Raven, 1997; Hanson et al., 2003; Engel et al., 2015; Fei et al., 2022; Hennacy et al., 2024) and is surrounded by a starch sheath that limits CO₂ escape (Toyokawa et al., 2020; He et al., 2023; Fig. 1a).

**Figure 1.**
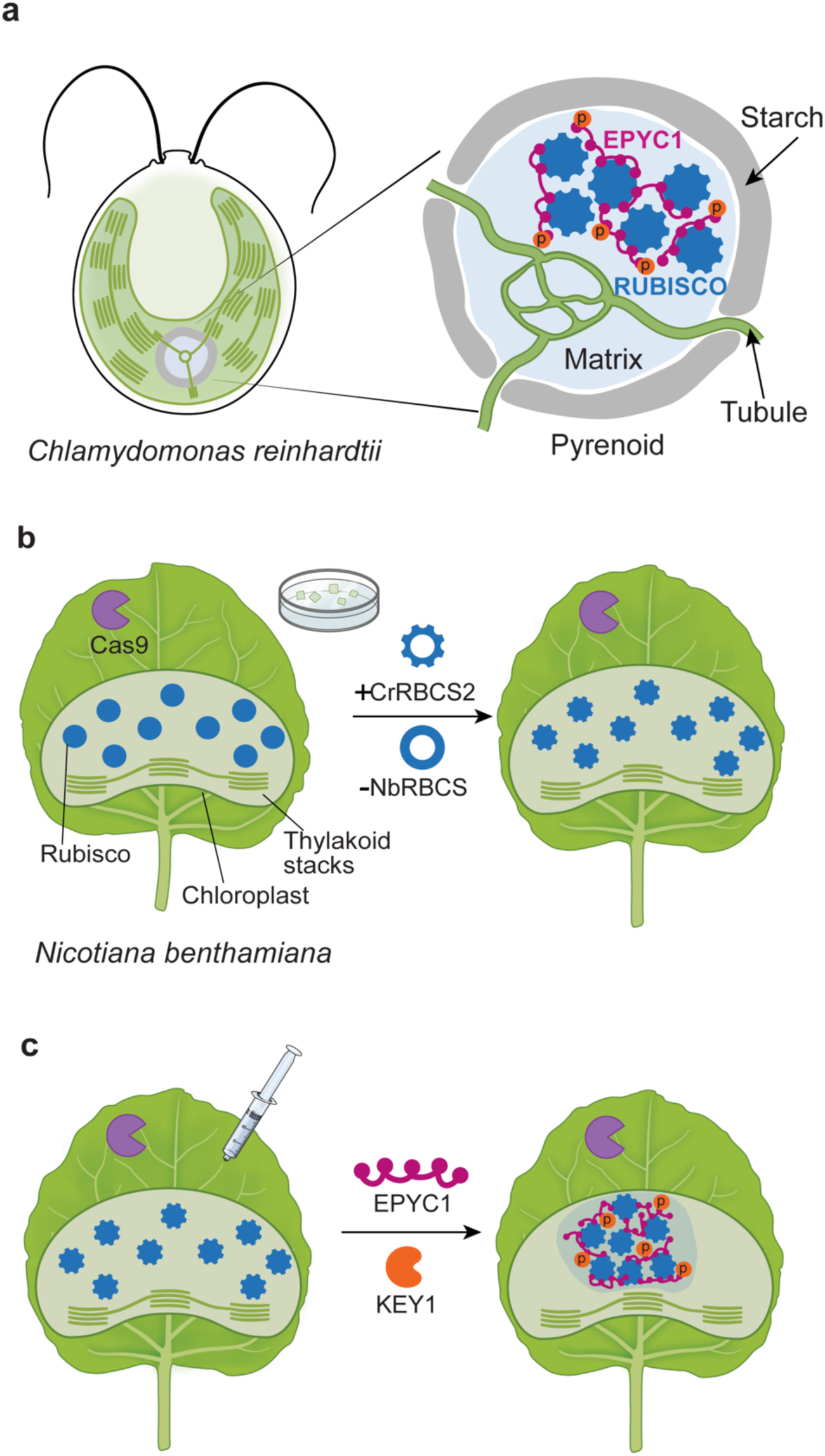
Overview of the *Chlamydomonas reinhardtii* pyrenoid and experimental strategy for reconstituting pyrenoid components in *Nicotiana benthamiana*. **(a)** The *C. reinhardtii* pyrenoid contains a liquid-like condensate of densely packed Rubisco and the intrinsically disordered protein EPYC1. The matrix is traversed by thylakoid tubules and surrounded by a starch sheath. Pyrenoid condensate number is regulated by the kinase KEY1, which phosphorylates EPYC1. **(b)** Replacement of endogenous *N. benthamiana* RBCS with *C. reinhardtii* RBCS2 (CrRBCS2) generates a Rubisco holoenzyme containing the *C. reinhardtii* small subunit. **(c)** Transient co-expression of EPYC1-GFP and KEY1-mScarlet in *Nbrbcs;CrRBCS2* plants results in consolidation of multiple condensates into a predominantly single condensate per chloroplast.

Considerable progress has been made toward reconstituting elements of the *C. reinhardtii* pyrenoid in plants, including the assembly of Rubisco condensates (Atkinson et al., 2020), partial reconstruction of matrix-traversing membranes (Hennacy et al., 2024), and recruitment of starch granules to the matrix (Atkinson et al., 2024). However, progress remains limited by an incomplete understanding of which components are required for minimal functional pyrenoid formation (Adler et al., 2022).

A major challenge in identifying these missing factors has been the relatively slow pace of testing candidates for assembly, localization, and regulation in an engineered system (Hennacy and Jonikas, 2020). *Arabidopsis thaliana* has served as a primary platform for stable expression of algal CCM components, but its limited amenability to transient expression limits effective iteration times to 4-6 months, imposed by transgenic plant generation time (Atkinson et al., 2017; Atkinson et al., 2020; Hennacy et al., 2024). Transgenerational protein silencing has further constrained systematic pyrenoid protein testing.

To address these bottlenecks of pyrenoid engineering in plants, here we sought to develop a Rubisco-recombinant system for testing pyrenoid components in the model plant *Nicotiana benthamiana*. In contrast to Arabidopsis, *N. benthamiana* has been widely used for transient protein expression, including in reconstitution of complex metabolic pathways (Lau and Sattely, 2015; Klein and Sattely, 2017; Bally et al., 2018; Dudley et al., 2022), and provides an effective system for testing protein localization *in planta* (Kazachkova et al., 2021). However, wild-type *N. benthamiana* had previously been of limited use in pyrenoid reconstitution because the endogenous Rubisco small subunit (RBCS) in *N. benthamiana* lacks the *C. reinhardtii*-specific residues required for binding of many pyrenoid proteins (Meyer et al., 2012; He et al., 2020; Meyer et al., 2020). The unmodified plant was therefore not expected to support pyrenoid assembly. Replacing endogenous RBCS with an algal counterpart is not straightforward: flowering plants encode large, multicopy *RBCS* gene families, and most loci must be disrupted to suppress competition from incompatible endogenous subunits.

In the present work, we produce a pyrenoid-compatible *N. benthamiana* by disrupting the endogenous *RBCS* genes and expressing their *C. reinhardtii* counterpart (Fig. 1b). We demonstrate the value of this system by reconstituting condensate formation using rapid, transient expression of EPYC1. We further use this system to establish that the number of pyrenoid condensates can be regulated in plants by expressing the pyrenoid kinase KEY1 (Fig. 1c). These results establish engineered *N. benthamiana* as a plant platform for rapid screening of candidate pyrenoid proteins and testing their roles in pyrenoid assembly, function, and regulation.

## Results

### CRISPR/Cas9-mediated genome editing disrupts RBCS accumulation in *N. benthamiana*

As a first step to establish a pyrenoid reconstitution system in an *N. benthamiana*, we needed remove and replace endogenous *NbRBCS* with the *C. reinhardtii* version to enable binding of *C. reinhardtii* pyrenoid components (Meyer et al., 2020). We queried the *N. benthamiana* genome assembly (Wang et al., 2024), accessed via Phytozome, https://phytozome-next.jgi.doe.gov with known *A. thaliana* RBCS protein sequences and identified eight photosynthetic *RBCS* loci, distributed across chromosomes 3, 17, 18, and 19. The eight loci fell into two structural groups based on exon organization: Group I (four exons, three introns) and Group II (three exons, two introns) (Supplementary Fig. 1a), consistent with the two *RBCS* subfamilies previously described in *Nicotianeae* (Jamet et al., 1991). One additional locus (Nbe11g19750) showed markedly lower sequence identity to the photosynthetic family (∼53% identity) and was excluded from targeting on the basis of its high sequence similarity to the trichome-specific NtRBCS-T protein described in tobacco (Laterre et al., 2017). We designed four sgRNAs to target conserved *RBCS* regions, ensuring that each of the eight loci was covered by at least two independent guides (Supplementary Fig. 1b).

We initially attempted *Agrobacterium*-mediated stable transformation with CRISPR-Cas9 overexpression constructs carrying single guide RNAs (sgRNAs), analogously to a recent approach used to disrupt *RBCS* in the related diploid species *Nicotiana sylvestris* (da Silva et al., 2026), but this approach yielded severely chlorotic T0 regenerants that failed to root and did not survive tissue culture even on sucrose-supplemented medium, precluding downstream complementation (Supplementary Fig. 2). We therefore employed a tissue culture-independent approach.

As a first step, we disrupted endogenous RBCS accumulation using CRISPR-Cas9-mediated genome editing with mobile sgRNAs (Ellison et al., 2020, Fig. 2a). This *in planta* editing approach relies on systemic delivery of sgRNAs via an RNA virus applied to the leaves of Cas9-expressing *N. benthamiana*, enabling multiplexed targeting of endogenous loci. Throughout this study, the wild-type control refers to this parental *N. benthamiana* line stably expressing Cas9 but lacking guide RNAs.

**Figure 2.**
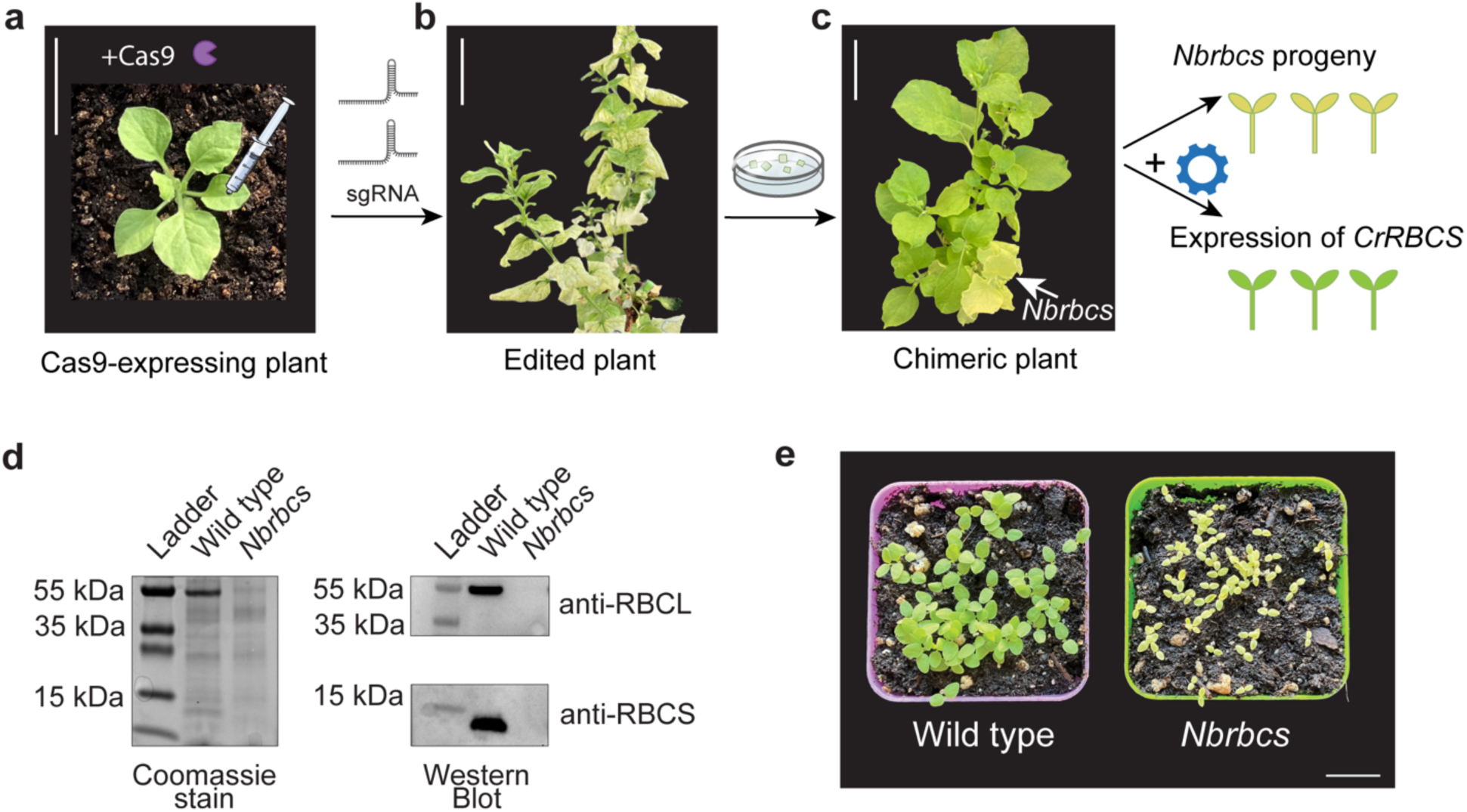
CRISPR-Cas9-mediated disruption of endogenous *NbRBCS* genes in *Nicotiana benthamiana* results in loss of autotrophic growth. **(a)** Experimental strategy for generating *Nbrbcs* plants. Cas9-expressing *N. benthamiana* plant prior to sgRNA delivery. **(b)** Edited plant two months after agroinfiltration of mobile sgRNAs targeting conserved regions of endogenous *NbRBCS* genes, showing white patchy leaves. **(c)** Chimeric plant regenerated from white leaf sections of the edited plant, showing white and green stems. White leaves of the chimeric plant were used for protein extraction and for *Agrobacterium*-mediated transformation with *CrRBCS2* to generate *Nbrbcs;CrRBCS2* lines. White stem of the chimeric plant produced viable seeds. Scale bars, 5 cm. **(d)** Immunoblot analysis of total protein extracted from wild-type and *Nbrbcs* leaf tissue. Membranes were probed with antibodies against NbRBCS and NbRbcL. Coomassie-stained gel is shown as a loading control. **(e)** Wild-type and *Nbrbcs* seedlings grown under greenhouse conditions. Scale bar, 2 cm.

Two months after infiltration with mobile sgRNAs, plants developed distinct white patches as well as fully white leaves, consistent with loss of endogenous *N. benthamiana* Rubisco and chlorophyll (Fig. 2b). This phenotype is characteristic of severe impairment of photosynthetic ability. We excised the white tissue and used it for full plant regeneration. Due to the severity of the phenotype, regenerated plants were not viable under greenhouse conditions, except for a single plant that exhibited chimeric white and green stems (Fig. 2c). The green stem likely retained functional Rubisco and supported growth of the adjacent white stem, whose bleached phenotype suggested a possible lack of Rubisco.

Leaves from the white stem (labeled *Nbrbcs*) of this chimeric plant were white and showed no detectable accumulation of endogenous NbRBCS or NbRbcL protein, as assessed by Western blot analysis (Fig. 2d). Leaf tissue from *N. benthamiana* wild-type plants analyzed in parallel showed robust accumulation of both NbRBCS and NbRbcL. The concomitant loss of NbRbcL in the white leaves of the chimeric plant is consistent with destabilization of the Rubisco holoenzyme in the absence of the NbRBCS small subunit. Progeny derived from the white part of the plant were uniformly white and failed to grow beyond the cotyledon stage when grown in the greenhouse (Fig. 2e), demonstrating stable disruption of endogenous NbRBCS protein accumulation following CRISPR-Cas9-mediated genome editing in *N. benthamiana*.

### Chlamydomonas reinhardtii RBCS complements Nbrbcs and supports autotrophic growth in N. benthamiana

To produce *N. benthamiana* plants compatible with *C. reinhardtii* pyrenoid components, we next expressed *C. reinhardtii Rubisco small subunit 2* (*CrRBCS2*; Cre02.g120150) in the *Nbrbcs* background and assessed its ability to support autotrophic growth (Fig. 3a). We obtained three independent transgenic *N. benthamiana* lines expressing *CrRBCS2* and we observed that *Nbrbcs;CrRBCS2-1,-2 and-3* complemented plants had restored green coloration as well as growth in the greenhouse (Fig. 3b).

**Figure 3.**
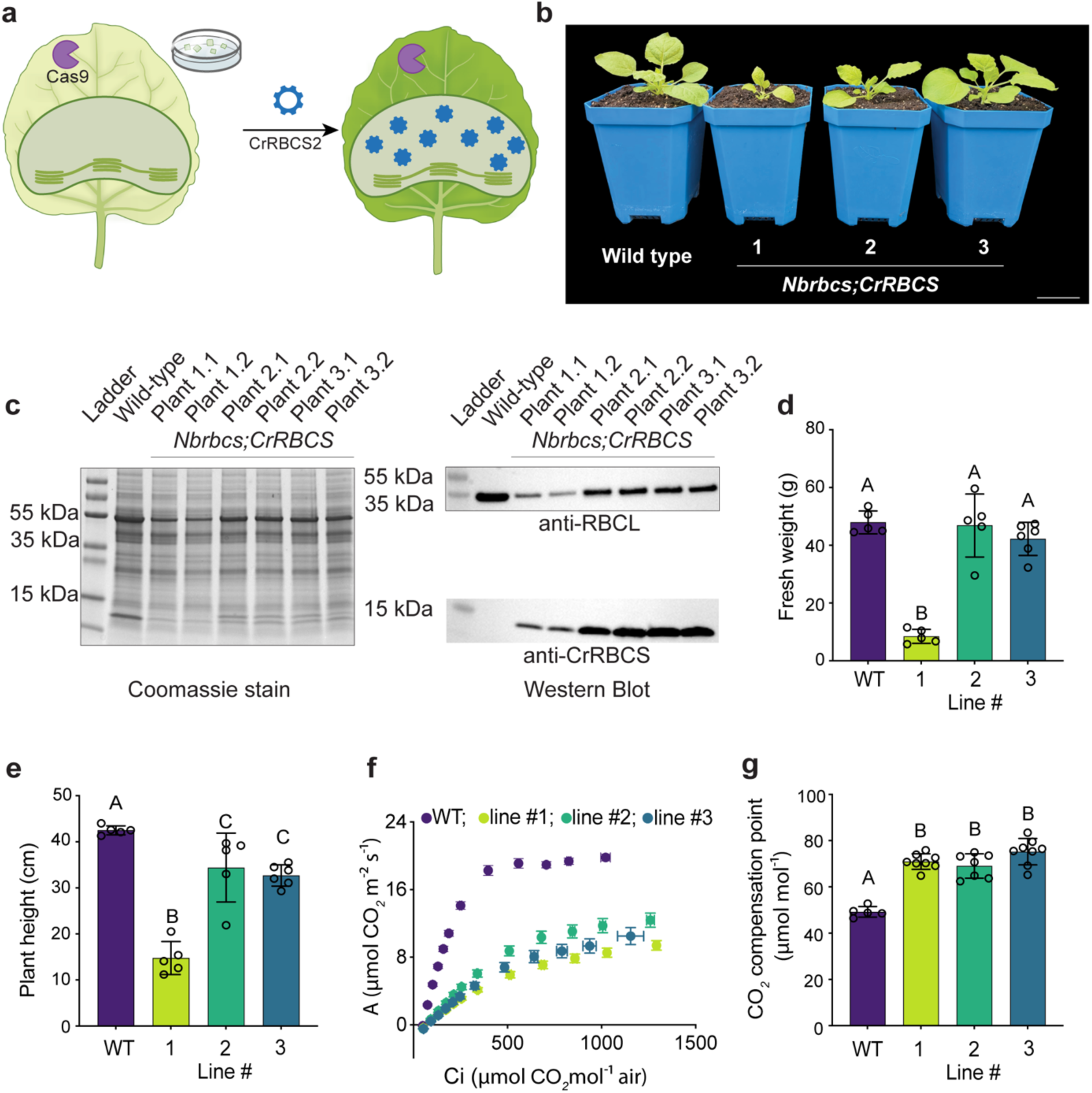
Complementation of *Nbrbcs* plants with *CrRBCS2* restores autotrophic growth. (a) Experimental strategy for complementation of *Nbrbcs* plants with *C. reinhardtii RBCS2* (*CrRBCS2*), generating the *Nbrbcs;CrRBCS2* line. (b) Representative photographs of wild-type and *Nbrbcs;CrRBCS2* lines grown under greenhouse conditions. Left to right: wild-type, Line 1, Line 2, Line 3. Scale bar, 5 cm. (c) Coomassie-stained gel and immunoblot analysis of total protein from wild-type and *Nbrbcs;CrRBCS2* lines. Membranes were probed with anti-CrRBCS and anti-NbRbcL antibodies. Coomassie-stained gel is shown as a loading control. (d) Fresh weight of wild-type and *Nbrbcs;CrRBCS2* lines. Data are mean ± SD. Different letters indicate statistically significant differences (Dunnett’s T3 post-hoc test). (e) Plant height of wild-type and *Nbrbcs;CrRBCS2* lines. Data are mean ± SD. Different letters indicate statistically significant differences (Dunnett’s T3 post-hoc test). (f) *A*/*C_i_* curves showing net CO₂ assimilation rate (*A*; µmol CO₂ m⁻² s⁻¹) as a function of intercellular CO₂ concentration (*C_i_*; µmol CO₂ mol⁻¹ air) for wild-type and *Nbrbcs;CrRBCS2* lines. Data are mean ± SE. (g) CO₂ compensation point of wild-type and *Nbrbcs;CrRBCS2* lines. Data are mean ± SD. Different letters indicate statistically significant differences (Dunnett’s T3 post-hoc test).

Western blot analysis of T1 progeny using a CrRBCS-specific antibody showed detectable accumulation of CrRBCS2, accompanied by restored accumulation of NbRbcL (Fig. 3c), suggesting that hybrid CrRBCS2/NbRbcL Rubisco holoenzymes could assemble in these plants.

We compared growth of the complemented lines with that of wild-type plants (Fig. 3d,e). Lines 2 and 3 showed fresh weight comparable to wild type but reduced plant height, whereas Line 1 showed the most severe reduction in both fresh weight and plant height (Fig. 3d,e). The more pronounced growth reduction in Line 1 may reflect lower Rubisco levels in this line. All the lines produced viable seeds, indicating that complementation restored viability across all three independent lines.

Leaf gas exchange measurements showed that all three *Nbrbcs;CrRBCS2* lines exhibited significantly lower net CO₂ assimilation rates than wild-type plants (Fig. 3f,g). Apparent *V_cmax_* values derived from *A*/*C_i_* curve fitting were substantially decreased, to 22%, 33%, and 29% of wild type in Lines 1, 2, and 3, respectively (Supplementary Table 1). These differences may reflect lower Rubisco expression in the transformant lines, altered catalytic performance of the hybrid enzyme, or both. The CO₂ compensation point was elevated in all three complemented lines; whether this reflects altered Rubisco oxygenase/carboxylase balance in the hybrid CrRBCS2/NbRbcL enzyme, reduced Rubisco abundance, or both, remains to be determined.

Stomatal conductance was comparable across all genotypes (Supplementary Table 1), indicating that reduced CO₂ assimilation was not attributable to stomatal limitation. Together with the reduced RbcL accumulation observed by immunoblot analysis (Fig. 3c), these results indicate that CrRBCS2 supports assembly of a functional Rubisco complex sufficient for autotrophic growth but does not fully restore photosynthetic capacity to wild-type levels.

Despite their reduced photosynthetic performance, Lines 2 and 3 displayed only modest growth defects under greenhouse conditions (Fig. 3d,e), likely reflecting the non-limiting growth conditions used in this study. We conclude that the Rubisco small subunit of *N. benthamiana* can be functionally replaced by that of *Chlamydomonas reinhardtii*.

### Transient expression of EPYC1 reconstitutes EPYC1-Rubisco matrix-like condensates in Rubisco-recombinant *N. benthamiana*

With a viable Rubisco-recombinant plant in hand, we next asked whether it supports algal pyrenoid matrix assembly. Matrix assembly in *C. reinhardtii* relies on multivalent interactions between Rubisco and the intrinsically disordered linker protein EPYC1 (He et al., 2020). To test whether transient EPYC1 expression is sufficient to promote condensate formation in our Rubisco-recombinant *N. benthamiana*, we transiently expressed EPYC1 as a GFP fusion under a constitutive promoter. Since it was previously shown that EPYC1 requires a chloroplast targeting signal for localization in *A. thaliana* (Atkinson et al., 2020) we also fused the *A. thaliana* RBCS1a chloroplast transit peptide to the N-terminus of EPYC1 (Fig. 4a).

**Figure 4.**
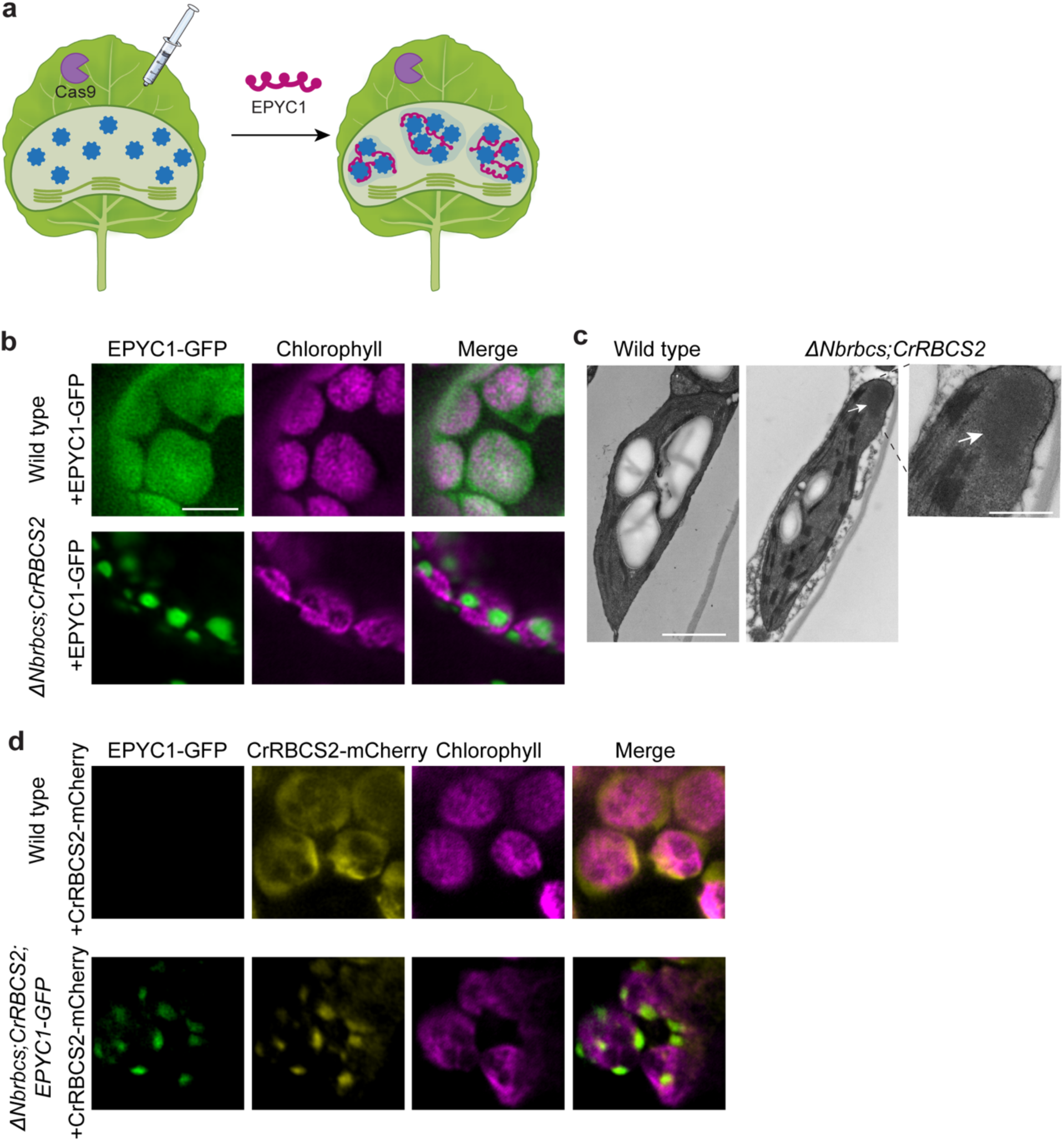
Transient expression of EPYC1-GFP in *Nbrbcs;CrRBCS2* chloroplasts results in formation of discrete condensates. (a) Experimental strategy for transient expression of EPYC1-GFP in *Nbrbcs;CrRBCS2* plants. (b) Confocal images of EPYC1-GFP localization in wild-type and *Nbrbcs;CrRBCS2* chloroplasts. Chlorophyll autofluorescence is shown in magenta. Scale bar, 5 µm. (c) Transmission electron microscopy of wild-type and *Nbrbcs;CrRBCS2;EPYC1-GFP* chloroplasts. Arrows indicate electron-dense structures within *Nbrbcs;CrRBCS2* chloroplasts. Scale bars, 1 µm (overview) and 500 nm (inset). (d) Confocal images of CrRBCS-mCherry localization in wild-type and *Nbrbcs;CrRBCS2;EPYC1-GFP* chloroplasts. Chlorophyll autofluorescence is shown in magenta. Scale bar, 5 µm.

We observed that transient expression of EPYC1-GFP in the *Nbrbcs;CrRBCS2* background consistently led to the formation of multiple discrete condensates of varying sizes within chloroplasts of all the three Rubisco-recombinant lines (Fig. 4b, Supplementary Fig. 3a). The condensates were also observed by transmission electron microscopy as distinct electron-dense structures within chloroplasts (Fig. 4c). In contrast, transient expression of EPYC1-GFP in wild-type *N. benthamiana* only resulted in diffuse stromal localization without detectable condensate formation (Fig. 4b), indicating that condensate formation depends on the presence of the *C. reinhardtii* Rubisco small subunit, as previously observed in *A. thaliana* (Atkinson et al., 2020). EPYC1 expression also did not cause visible leaf phenotypes such as chlorosis or necrosis in either wild-type or recombinant contexts, suggesting little to no toxic effects of exogenous expression.

To rule out the possibility that the multiple condensates we observed in the *Nbrbcs;CrRBCS2* plants are an artifact of transient expression, we generated stable *EPYC1*-*GFP*-expressing *Nbrbcs;CrRBCS2* lines in the *Nbrbcs;CrRBCS2-2* background. We obtained two independent transgenic lines, both of which exhibited a multiple-condensate phenotype similar to that observed upon transient expression, indicating that multiple condensate formation is reproducible across expression modes (Supplementary Fig. 3b,c). We conclude that transient or stable expression of EPYC1 in Rubisco-recombinant *N. benthamiana* results in multiple pyrenoid matrix-like condensates.

The pyrenoid matrix forms by the co-condensation of Rubisco and EPYC1 (Mackinder et al., 2016; Wunder et al., 2018), so to confirm that the condensates we observed actually represent a reconstituted matrix, we transiently expressed CrRBCS2-mCherry in stable *EPYC1*-*GFP*-expressing *Nbrbcs;CrRBCS2* lines. CrRBCS2-mCherry signal co-localized with EPYC1-GFP condensates (Fig. 4d), demonstrating that Rubisco is enriched within the condensates. Together with the strict dependence of condensate formation on the CrRBCS2-containing background, these data establish that EPYC1 has phase-separated with CrRBCS2-containing Rubisco, reconstituting in *N. benthamiana* the core compositional features of the *C. reinhardtii* pyrenoid matrix.

### Pyrenoid kinase KEY1 promotes single condensate formation in *N. benthamiana* chloroplasts

In *C. reinhardtii,* the pyrenoid kinase KEY1 is required to maintain a single pyrenoid per chloroplast (He et al., 2026). KEY1 is thought to promote exchange of Rubisco and EPYC1 with the stroma, which results in dissolution of smaller condensates and growth of a single larger one through Ostwald ripening. The loss of KEY1 results in multiple pyrenoid condensates, and such mutants show impaired growth under ambient CO₂ conditions, suggesting that consolidation into a single condensate may be important for efficient CO₂ concentration around Rubisco (He et al., 2026).

Considering that the multiple-condensate phenotype we observed in our *N. benthamiana* system (Fig. 4, Supplementary Fig. 3b,c) resembled the multiple-condensate phenotype of *C. reinhardtii key1* mutants, we reasoned that expression of KEY1 may promote a single condensate.

To test whether KEY1 can regulate condensate number in a heterologous plant system, we expressed full-length, codon-optimized *C. reinhardtii KEY1* fused to *mScarlet3* in *Nbrbcs;CrRBCS2-2* plants (Fig. 5a). Without a chloroplast transit peptide, KEY1-mScarlet localized to the cytosol and nucleus (Supplementary Fig. 3d). Adding a chloroplast transit peptide redirected KEY1-mScarlet to the chloroplast stroma (Fig. 5b). Co-expressing chloroplast-targeted *KEY1* with *EPYC1-GFP* markedly shifted condensate organization. Whereas EPYC1-GFP typically formed multiple condensates per chloroplast in the *Nbrbcs;CrRBCS2-2* background, co-expressing it with KEY1 predominantly produced a single condensate per chloroplast (Fig. 5b,c). Quantifying the condensates per chloroplast revealed a significant decrease in condensate number upon KEY1 co-expression (EPYC1-GFP: median 4, n = 130; EPYC1-GFP + KEY1-mScarlet: median 1, n = 153; Mann-Whitney U test, p < 0.0001; Fig. 5d). We consistently observed this change in condensate number across three independent experiments.

**Figure 5.**
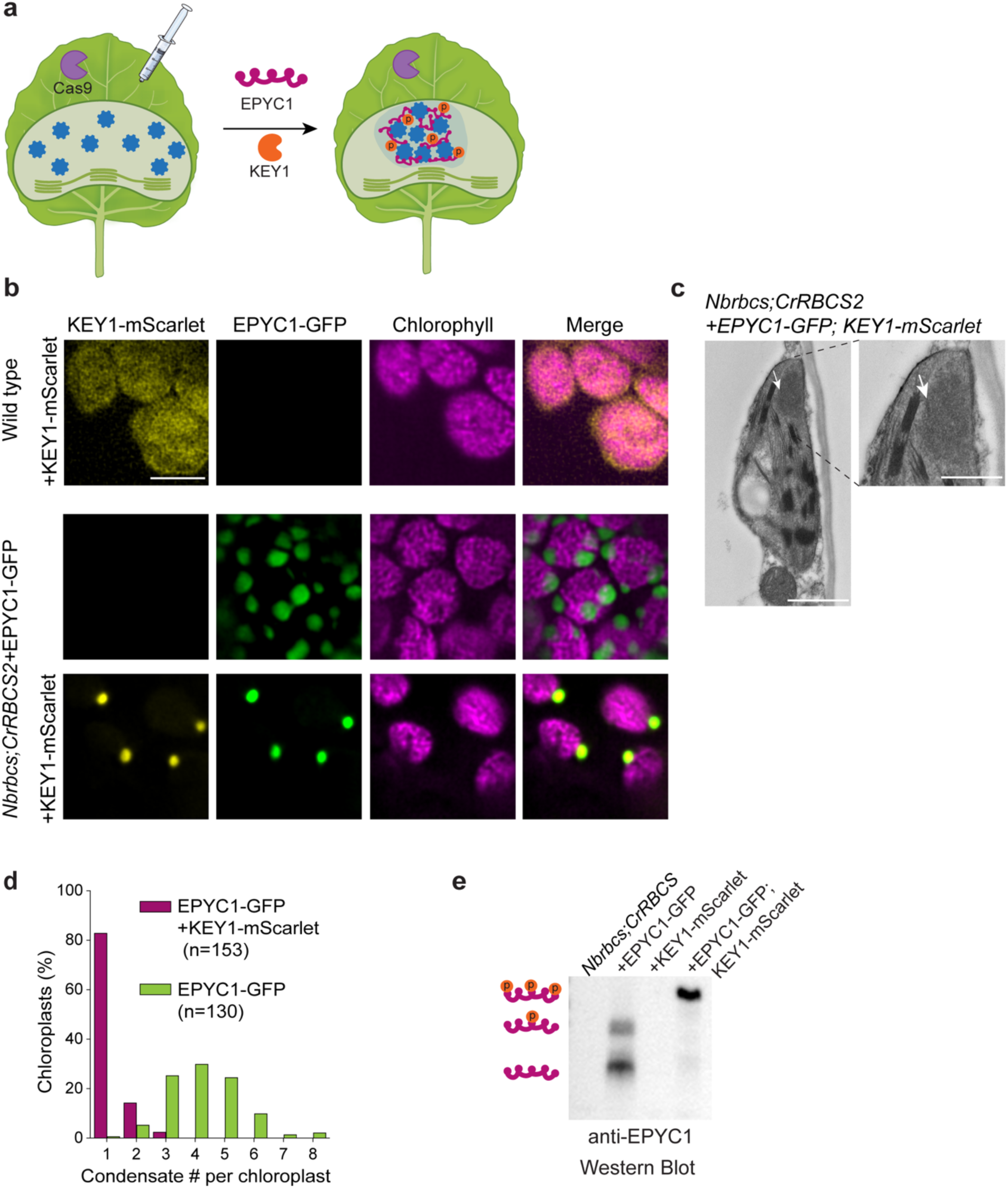
Co-expression of KEY1-mScarlet and EPYC1-GFP in *Nbrbcs;CrRBCS2* chloroplasts results in a single condensate per chloroplast. (a) Experimental strategy for co-expression of KEY1-mScarlet and EPYC1-GFP in *Nbrbcs;CrRBCS2* plants. (b) Confocal images of KEY1-mScarlet and EPYC1-GFP localization in wild-type and *Nbrbcs;CrRBCS2* chloroplasts. Chlorophyll autofluorescence is shown in magenta. Scale bar, 5 µm. (c) Distribution of condensate number per chloroplast in *Nbrbcs;CrRBCS2* plants expressing EPYC1-GFP alone or co-expressing EPYC1-GFP and KEY1-mScarlet. n = 130 chloroplasts (EPYC1-GFP) and n = 153 chloroplasts (EPYC1-GFP + KEY1-mScarlet) from three independent experiments. Mann-Whitney U test, p < 0.0001. (d) Transmission electron microscopy of *Nbrbcs;CrRBCS2*+EPYC1-GFP; KEY1-mScarlet chloroplasts. Arrows indicate electron-dense structures. Scale bars, 1 µm (overview) and 500 nm (inset). (e) Immunoblot analysis with anti-EPYC1 antibody following Phos-tag SDS-PAGE of protein extracted from *Nbrbcs;CrRBCS2* plants expressing EPYC1-GFP alone, KEY1-mScarlet alone, or EPYC1-GFP and KEY1-mScarlet together.

KEY1 phosphorylates EPYC1 *in vitro* and is required for EPYC1 phosphorylation in *C. reinhardtii* (He et al., 2026). To examine whether KEY1 expression alters EPYC1 phosphorylation status, total protein extracts from plants expressing EPYC1-GFP alone or together with KEY1-mScarlet were analyzed by Phos-tag SDS–PAGE and immunoblotting with anti-EPYC1 antibodies. In the presence of KEY1, EPYC1 consistently exhibited decreased electrophoretic mobility relative to EPYC1 expressed alone (Fig. 5e). These data suggest that KEY1 phosphorylates EPYC1 *in planta*, resulting in a reproducible change in EPYC1 mobility on Phos-tag SDS-PAGE gels. Together, these results indicate that KEY1 promotes consolidation of multiple Rubisco-EPYC1 condensates into a single condensate per chloroplast, apparently through phosphorylation of EPYC1 *in planta*. These findings demonstrate that the core regulatory logic controlling pyrenoid number in algae is portable and functional in a heterologous plant system.

## Discussion

Here we report the generation of a Rubisco-recombinant *N. benthamiana* platform that enables rapid functional characterization of pyrenoid components in a plant context. By replacing endogenous *NbRBCS* loci with the *C. reinhardtii* counterpart, we obtained plants expressing a hybrid plant-algal Rubisco holoenzyme compatible with algal pyrenoid proteins. Transient expression of EPYC1 in this background drove Rubisco-EPYC1 condensate formation, and co-expression of the pyrenoid kinase KEY1 consolidated multiple condensates into a single condensate per chloroplast, apparently through phosphorylation of EPYC1 *in planta*.

The generation of the Rubisco-recombinant line required overcoming a fundamental obstacle for a photosynthetic organism: Rubisco is essential for photoautotrophic growth. In *N. benthamiana*, complete disruption of the eight *NbRBCS* loci through conventional stable transformation produced T0 regenerants that failed to survive tissue culture, precluding the stable transformation needed for complementation. We therefore used mobile sgRNA delivery via agroinfiltration into Cas9-expressing plants (Ellison et al., 2020) which produced edited plants with white and green mosaic leaves. Regeneration from white leaf tissue yielded, serendipitously, a single chimeric regenerant in which a green stem likely sustained adjacent Rubisco-null white tissue through intercellular metabolite exchange (Fig. 2c), an arrangement functionally analogous to deliberate grafting strategies used to sustain photosynthetically-impaired plants (Kanevski and Maliga, 1994). We used white tissue from this plant directly as a source for stable transformation with *CrRBCS2*, bypassing the lethality that would otherwise preclude recovery of the complemented line. The exploitation of a naturally occurring chimeric regenerant as a transformation intermediary has not, to our knowledge, been previously described, and may offer a broadly applicable strategy for engineering essential pathways in plants.

We demonstrate here that this Rubisco-recombinant background enabled condensate formation upon transient expression of GFP-tagged EPYC1, making our engineered *N. benthamiana* the second plant species after *A. thaliana* in which Rubisco condensates have been reconstituted by stable EPYC1 expression (Atkinson et al., 2020). This result suggests that Rubisco condensate formation can be robustly achieved across plant species when appropriate molecular interfaces are engineered (He et al., 2020; Meyer et al., 2020). However, in *N. benthamiana*, the expression of GFP-tagged EPYC1 consistently resulted in multiple condensates per chloroplast (Fig. 4, Supplementary Fig. 3a) rather than the single punctum typically observed in Arabidopsis (Atkinson et al., 2020). This phenotype was observed across both transient and stable expression contexts, suggesting that formation of multiple condensates represents a default outcome of EPYC1-driven phase separation in *N. benthamiana* chloroplasts (Supplementary Fig. 3b,c). One possible explanation for this difference is that the less-complete replacement of RBCS in Arabidopsis may result in weaker EPYC1-Rubisco interactions, which could promote Ostwald ripening toward a single dominant condensate. Alternatively, an endogenous chloroplast kinase in *A. thaliana* may phosphorylate EPYC1 and promote condensate ripening. This latter idea raises the more general possibility that different host plant species may influence pyrenoid assembly in different ways that will need to be accounted for in engineering efforts.

In algae, loss of the pyrenoid-associated kinase KEY1 is linked to the formation of multiple pyrenoid condensates, and KEY1 has been shown to control pyrenoid number through phosphorylation of EPYC1 (He et al., 2026). In the engineered *N. benthamiana* system, co-expression of KEY1 with EPYC1 reorganized the multi-condensate state into a predominantly single condensate per chloroplast, and KEY1 expression was associated with a reproducible shift in EPYC1 mobility on Phos-tag gels, consistent with the KEY1-dependent phosphorylation of EPYC1 *in planta* (Fig. 5). KEY1 expression alone did not induce condensate formation, supporting the idea that its role in this system is regulatory rather than structural. We note that in *C. reinhardtii* and *in vitro*, very high levels of EPYC1 phosphorylation led to full dissolution of EPYC1-Rubisco condensates (He et al., 2026); technical limitations of possible chlorophyll bleedthrough prevented us from reliably distinguishing cells with fully dissolved condensates, if they exist in our system, from cells not expressing KEY1. Together, these results demonstrate that regulation of condensate number can be layered onto a minimal EPYC1-Rubisco condensate and that key aspects of pyrenoid regulation can be reconstituted across the green lineage.

The *N. benthamiana* Rubisco-recombinant platform described here is designed for modularity and rapid component iteration. Transient expression of candidate pyrenoid proteins can be completed from cloning to results within 14 days, enabling rapid assessment of localization and interaction with other pyrenoid components. Proteins required for pyrenoid-traversing membrane formation (Hennacy et al., 2024), carbonic anhydrase localization to the pyrenoid lumen, and starch sheath assembly remain either unidentified or poorly characterized in a heterologous context and represent immediate targets for testing in this system. Stable EPYC1-GFP lines, in which Rubisco-EPYC1 condensates are constitutively present, provide a defined starting point for introducing additional candidate proteins by transient expression. This creates a branching framework where each stable expression milestone enables the next round of rapid screening. Together with the condensate reconstitution and regulatory results presented here, this stepwise approach, moving from transient validation to stable integration, lays the conceptual and experimental foundation for the systematic reconstitution of a functional pyrenoid in plants and, ultimately, in agriculturally relevant crop species.

## Materials and Methods

### Plant material and generation of transgenic plants

*N. benthamiana* plants were grown in a climate-controlled greenhouse at 24 °C with natural light. Seeds of *N. benthamiana* stably expressing Cas9 were obtained from the laboratory of Dr. Daniel Voytas (University of Minnesota, Twin Cities). To generate *NbRBCS* knockout plants, the pEE83 construct harboring four *NbRBCS*-specific sgRNAs was introduced into *Cas9*-expressing plants by *Agrobacterium*-mediated infiltration as described in Ellison et al. (2020). After 6–8 weeks, white leaves were excised, sterilized, and used for whole-plant regeneration (Clemente, 2006). A single chimeric plant exhibiting white and green stems was recovered from regeneration and transferred to the greenhouse. White leaves from the white stem of this chimeric plant were used for *Agrobacterium*-mediated stable transformation with a binary vector harboring *CrRBCS2* under the control of the *Arabidopsis thaliana RBCS1A* promoter. To generate stable EPYC1-expressing lines, the *Nbrbcs;CrRBCS2-2* line was transformed by *Agrobacterium*-mediated stable transformation with a binary vector harboring *EPYC1-GFP* under the control of the *CsVMV* promoter. Transgenic plants were selected on hygromycin (20 mg/L) or glufosinate ammonium (15 mg/L). Positive transformants were confirmed by confocal microscopy and immunoblotting.

Fresh weight and plant height were measured in 2.5-month-old *Nbrbcs;CrRBCS2* and wild-type plants grown in a greenhouse under natural light conditions. Above-ground biomass was harvested and weighed immediately. Plant height was measured from soil level to shoot tip. Five to six biological replicates per line were measured.

### Transient transformation of *N. benthamiana*

*Agrobacterium*-mediated transient gene expression in *N. benthamiana* was carried out according to Zhang et al., with minor modifications (Zhang et al., 2020). *Agrobacterium* strains carrying each construct were grown overnight at 28 °C on LB plates containing gentamicin (25 mg/L) and the appropriate selective antibiotic for each construct. On the day of infiltration, cultures were resuspended to OD₆₀₀=0.3 in infiltration medium (0.25× Murashige and Skoog, pH 6.0, 1% sucrose, 100 µM acetosyringone, 0.005% Silwet L-77) and incubated in the dark for 3 h. Four-week-old plants were infiltrated by syringe into fully expanded leaves 3–5, kept in the dark for 24 h at 24 °C, and imaged 4 days post-infiltration. Where indicated, multiple constructs were mixed at equal ratios prior to infiltration.

### Plant imaging

Four-mm leaf discs were cut using a biopsy punch, mounted in perfluoroperhydrophenanthrene on glass slides (Littlejohn et al., 2014). The slides were imaged using a VT-iSIM super-resolution confocal microscope on an Olympus iX83 body equipped with a Hamamatsu ORCA Quest sCMOS camera and running VisiView control software.

GFP was excited at 488 nm and collected at 545/50 nm; mScarlet and mCherry were excited at 561 nm and collected at 595/40 nm; chlorophyll autofluorescence was excited at 642 nm and collected at 700/75 nm. Z-series were acquired with 300 nm steps using a 60× oil-immersion objective, NA 1.42. Images were processed in Fiji using Microvolution deconvolution plugin (Schindelin et al., 2012). Images were acquired at 1×1 binning.

### Construct design and cloning

Constructs for EPYC1 and KEY1 expression were assembled using the GoldenBraid cloning system and GoldenBraid binary vectors obtained from Addgene (Sarrion-Perdigones et al., 2013). *EPYC1-GFP* was assembled as a fusion of the *Arabidopsis thaliana RBCS1A* chloroplast transit peptide at the N-terminus, followed by *EPYC1* and C-terminal *GFP*, under the control of the *CsVMV* promoter. *CrRBCS2-mCherry* was assembled as a fusion of the *Arabidopsis thaliana RBCS1A* chloroplast transit peptide at the N-terminus, followed by *CrRBCS2* and C-terminal *mCherry*, under the control of the tomato Ubiquitin 10 (*SlUBQ10*) promoter. KEY1-mScarlet3 was assembled in two versions: one carrying the *AtRBCS1A* chloroplast transit peptide at the N-terminus and one lacking a transit peptide, both under the control of the *SlUBQ10* promoter. Synthetic DNA fragments codon optimized for *Nicotiana benthamiana* expression were used for assembly. The *CrRBCS2* complementation construct, consisting of the *AtRBCS1A* promoter and transit peptide driving expression of *CrRBCS2*, was kindly provided by the McCormick laboratory (University of Edinburgh). Four sgRNAs targeting *Nicotiana benthamiana RBCS* genes were cloned into the pEE83 vector obtained from Addgene (deposited by the Daniel Voytas laboratory). All final constructs were confirmed by full plasmid sequencing and introduced into *Agrobacterium tumefaciens* GV3101 by electroporation.

### Photosynthetic measurements

Leaf gas exchange was measured using a LI-6800 Portable Photosynthesis System (LI-COR Biosciences) on leaves 3–5 of 2 month-old plants grown in a greenhouse under natural light conditions. Unless stated otherwise, chamber conditions were set to: T_xchg_ 23.4°C, reference CO₂ 440 µmol mol^−1^, PPFD 1500 µmol m^−2^ s^−1^, and airflow 500 µmol s^−1^, corresponding to an average VPD of 1.89 ± 0.27 kPa. Fully expanded leaves were clamped in the chamber, then net CO₂ assimilation (*A*), stomatal conductance (*g_s_*), and intercellular CO₂ (*C_i_*) were recorded at steady state. To establish steady-state stomatal conductance, leaves were first acclimated to a CO₂ level of 400 µmol mol^−1^. CO₂ was then stepped down to 50 µmol mol^−1^ and held until stability was reached, allowing stomata and Rubisco activation state to equilibrate before measurements began. These acclimation points were excluded from the final *A*/*C_i_* curve. *A*/*C_i_* curves were generated by stepping CO₂ from 50 to 1500 µmol mol^−1^ across 12 setpoints.

To determine the CO₂ compensation points, a linear regression was fit to the first four points on each *A*/*C_i_* curve. The compensation point (Γ) was the *C_i_* calculated from each regression by setting *A* equal to zero. The maximum rate of Rubisco carboxylation (*V_cmax_*) and maximum electron transport rate (*J_max_*) were estimated by fitting *A*/*C_i_* curves to the Farquhar–von Caemmerer–Berry (FvCB) model of C3 photosynthesis (Farquhar et al., 1980) using the R package plantecophys v1.4 (Duursma, 2015). Rubisco kinetic constants and their temperature dependencies were taken from Bernacchi et al. (2001). *g_s_* was recorded at the ambient CO₂ step (∼400 µmol mol^−1^). Because *V_cmax_* was calculated using kinetic parameters derived from tobacco Rubisco (Bernacchi et al., 2001), these values are comparative estimates of leaf-level CO₂ assimilation. Measurements were obtained from 5–7 biological replicates per genotype, one leaf per plant.

### Immunoblot analysis

Total soluble protein was extracted in buffer (20 mM Tris-HCl pH 7.5, 5 mM MgCl₂, 300 mM NaCl, 5 mM DTT, 1% Triton X-100, cOmplete Mini EDTA-free Protease Inhibitor Cocktail (Roche)). Samples were centrifuged at 16,000 × g for 10 min and the supernatant was heated at 80 °C for 10 min with 1× Laemmli Sample Buffer (Bio-Rad Laboratories) and 200 mM DTT. Extracts were separated on precast 4–20% gels (Bio-Rad Laboratories) and transferred to Immobilon-P PVDF (MilliporeSigma) by wet transfer at 100 V for 1 h at 4 °C. Membranes were blocked in 5% dry nonfat milk in TBST, incubated with primary antibody in 2.5% milk overnight at 4 °C, washed three times in TBST, and incubated with HRP-linked goat anti-rabbit IgG (Abcam; 1:10,000). Detection was performed using WesternBright ECL (Advansta). Membranes were probed with custom peptide antibodies against RbcL (1:5,000), CrRBCS (1:10,000) produced by YenZym Antibodies, LLC.

For Phos-tag SDS–PAGE, leaf tissue was homogenized in extraction buffer (5 mM HEPES-KOH pH 7.5, 100 mM DTT, 100 mM Na₂CO₃, 2% SDS, 12% sucrose, 1 mM NaF, 0.3 mM Na₃VO₄, 2× Halt Protease Inhibitor Cocktail EDTA-free (Thermo Fisher Scientific), and 2× Halt Phosphatase Inhibitor Single-Use Cocktail (Thermo Fisher Scientific)) and clarified by centrifugation at 16,000 × g for 10 min at 4 °C. Clarified extracts were heated at 70 °C for 10 min with 1× Laemmli Sample Buffer (Bio-Rad Laboratories) and 200 mM DTT. Samples were resolved on 12.5% SuperSep™ Phos-tag™ precast gels (Fujifilm) and electrophoresed at 100 V for 200 min. Proteins were transferred to Immobilon-P PVDF membranes (MilliporeSigma) by wet transfer at 100 V for 1 h at 4 °C. Following transfer, membranes were incubated with anti-EPYC1 antibody (YenZym Antibodies, LLC; 1:5,000 in TBST containing 2.5% non-fat dry milk), washed three times in TBST, and incubated with HRP-linked goat anti-rabbit IgG (Abcam; 1:10,000). Signals were detected using WesternBright ECL (Advansta) and visualized using an iBright FL1500 Imaging System (Thermo Fisher Scientific).

### Transmission electron microscopy

Samples were fixed in 2.5% glutaraldehyde (Sigma-Aldrich) and washed with PBS. Samples were dehydrated in a graded ethanol series (30–100%), transitioned through propylene oxide (Electron Microscopy Sciences), and embedded in LR White resin (Electron Microscopy Sciences). Sections (80 nm) were cut using an Ultramicrotome EM UC7 (Leica) and mounted on mesh grids. Grids were stained with 2% uranyl acetate (Electron Microscopy Sciences) followed by 1% lead citrate (Electron Microscopy Sciences) and imaged using a transmission electron microscope HT7800 (Hitachi) equipped with a Morada G3 digital camera (EMSIS).

## Acknowledgements

We thank Dr. Sabrina Ergun, Dr. Aastha Garde, Dr. Andrii Vainer, Marie Bao, as part of Life Science Editors, as well as all the members of the Jonikas lab for helpful discussions and feedback. Seeds of Cas9-expressing *N. benthamiana* plants were kindly provided by Dr. Evan Ellison and Dr. Daniel Voytas (University of Minnesota). We thank Dr. Nicky Atkinson and Dr. Alistair McCormick (University of Edinburgh) for advice on the project and for providing the *CrRbcS2* expression plasmid. This project was supported by grants from the National Institute of General Medical Sciences of the National Institutes of Health (T32GM007388); the US National Science Foundation (MCB-1935444), the US National Institutes of Health (1R01GM140032-01), the Howard Hughes Medical Institute, the Carbon Technology Research Foundation (AP23-1_023), and the Bill & Melinda Gates Foundation and United Kingdom Foreign, Commonwealth and Development Office (INV-054558) to M.C.J. The work conducted by the US Department of Energy Joint Genome Institute (https://ror.org/04xm1d337), a DOE Office of Science User Facility, was supported by the Office of Science of the US Department of Energy under Contract No. DE-AC02-05CH11231 (https://doi.org/10.46936/10.25585/60008814). The content is solely the responsibility of the authors and does not necessarily represent the official views of the funders.

## Author Contributions

Y.K. and M.C.J. conceived the study. Y.K. and J.I.P. performed the experiments. A.K.R. contributed to experimental design and to establishing and optimizing LI-COR gas exchange measurements. E.F. contributed to transmission electron microscopy. Y.K. and M.C.J. wrote the manuscript with input from all authors.

---

**Supplementary Figure 1.**
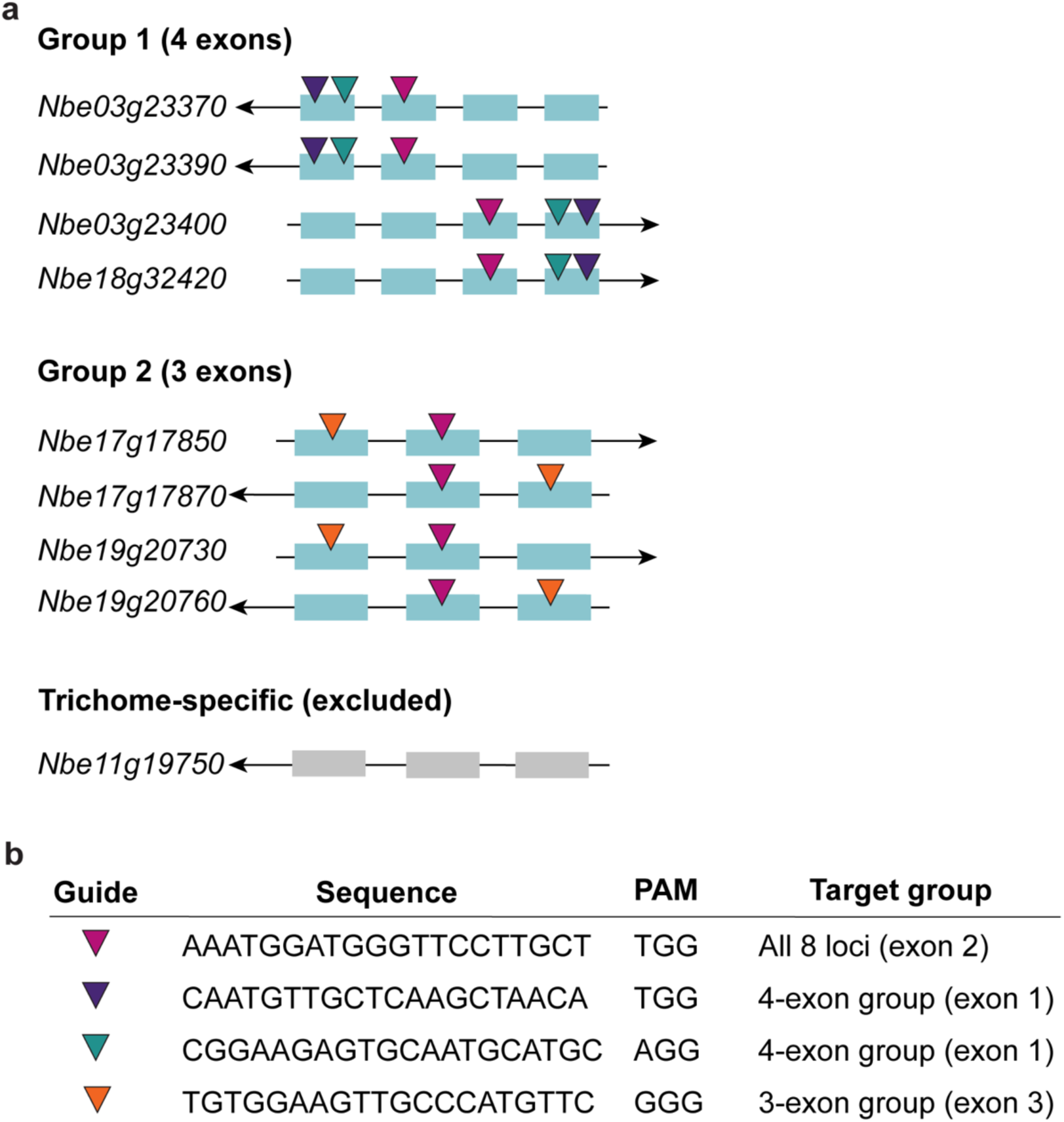
Gene structure and sgRNA target sites of photosynthetic *NbRBCS* loci in *N. benthamiana*. (a) Gene models of the eight photosynthetic *NbRBCS* loci identified in the *N. benthamiana* genome assembly. Loci are grouped by exon number: Group I (4 exons) and Group II (3 exons). Arrows indicate transcription orientation. Colored triangles indicate sgRNA target positions. One trichome-specific locus (Nbe11g19750) is shown in gray and was excluded from targeting. (b) Sequences of the four sgRNAs used for *NbRBCS* disruption, with corresponding PAM sequences and target groups.

**Supplementary Figure 2.**
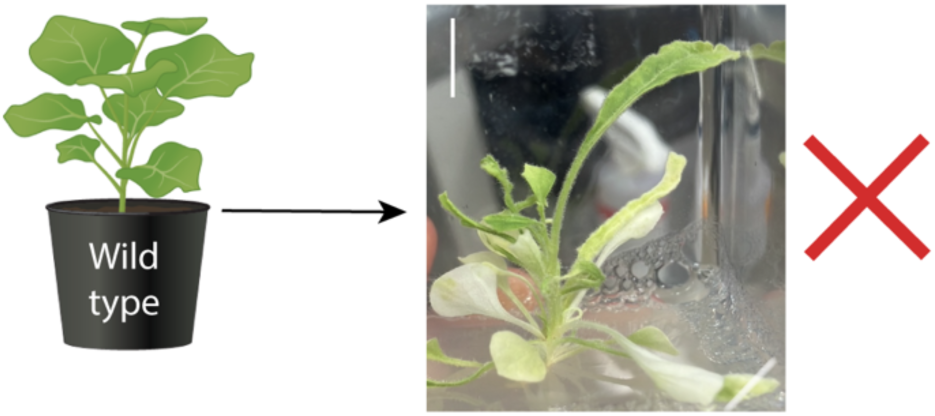
Stable transformation with a Cas9/sgRNA construct targeting *NbRBCS* genes does not yield viable *Nbrbcs* plants. Representative T0 regenerant following stable transformation with a Cas9/sgRNA construct targeting *NbRBCS* genes. T0 regenerants failed to root on sucrose-supplemented Murashige and Skoog (MS) medium and died before completing a full life cycle. Complementation with *CrRBCS2* was unsuccessful. Scale bar, 1 cm.

**Supplementary Figure 3.**
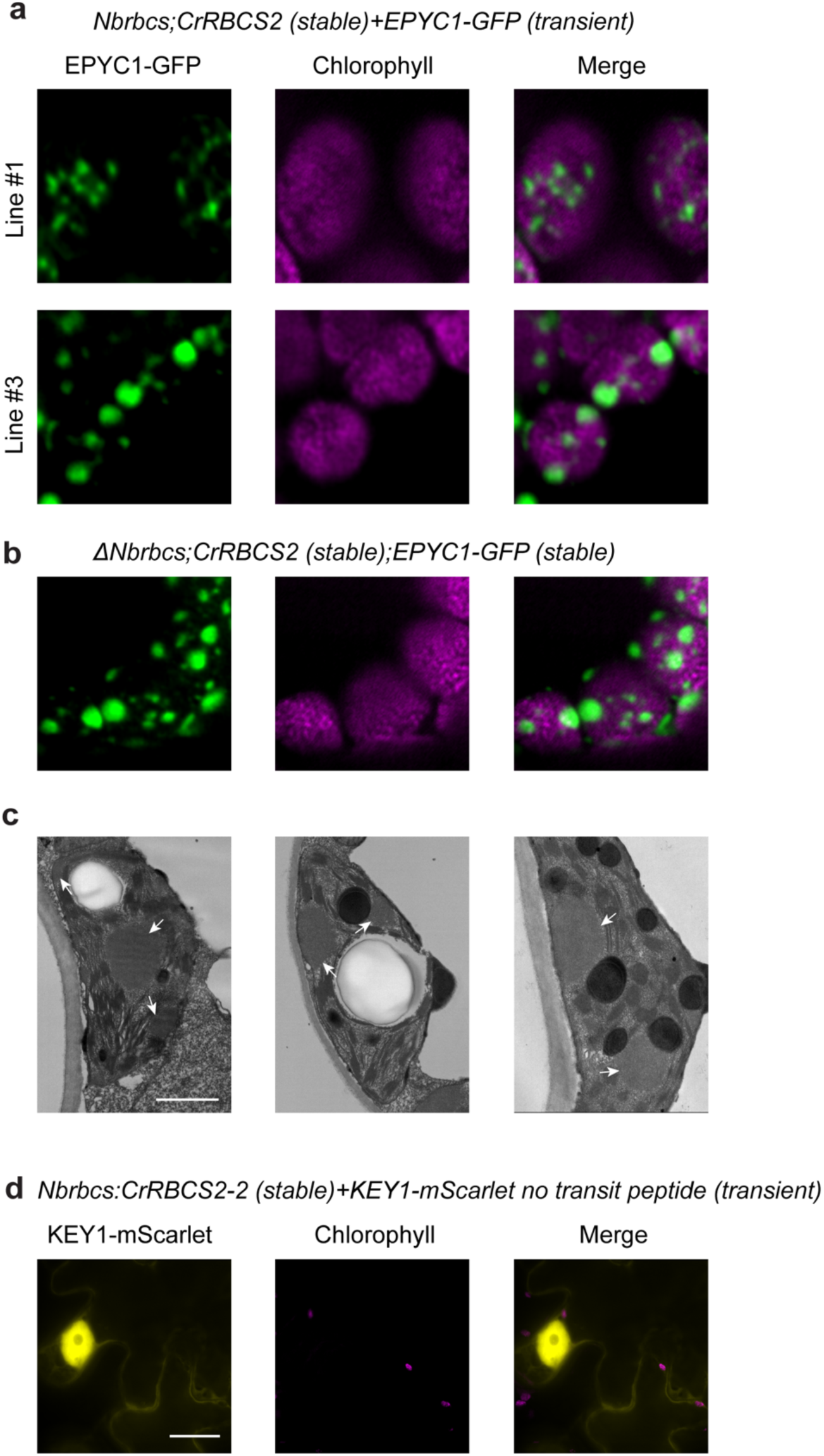
Additional characterization of EPYC1-GFP condensates and KEY1-mScarlet localization in *Nbrbcs;CrRBCS2* chloroplasts. (a) Confocal images of EPYC1-GFP localization in *Nbrbcs;CrRBCS2* Line 1 and Line 3 following transient expression. Chlorophyll autofluorescence is shown in magenta. Scale bar, 5 µm. (b) Confocal images of EPYC1-GFP localization in *Nbrbcs;CrRBCS2* plants following stable expression. Chlorophyll autofluorescence is shown in magenta. Scale bar, 5 µm. (c) Transmission electron microscopy of *Nbrbcs;CrRBCS2;EPYC1-GFP* chloroplasts. Arrows indicate electron-dense structures. Scale bars, 1 µm (overview) and 500 nm (insets). (d) Confocal images of KEY1-mScarlet expressed without a chloroplast transit peptide in *Nbrbcs;CrRBCS2* plants. Chlorophyll autofluorescence is shown in magenta. Scale bar, 5 µm.

**Supplementary Table 1.** Photosynthetic parameters of *Nbrbcs;CrRBCS2* complemented lines and wild-type *N. benthamiana*. *V_cmax_*, maximum rate of Rubisco carboxylation; *J_max_*, maximum rate of electron transport; *G_s_*, stomatal conductance at ambient CO₂. Parameters estimated by fitting *A/C_i_* curves to the Farquhar–von Caemmerer–Berry model (Farquhar et al., 1980) using kinetic constants from Bernacchi et al. (2001). Values are mean ± SD. *G_s_* measured at ambient CO₂ (∼400 µmol mol⁻¹).

| Genotype | n | $V_{cmax}$ mean ± SD<br>($\mu\text{mol CO}_2 \text{ m}^{-2} \text{ s}^{-1}$) | $J_{max}$ mean ± SD<br>($\mu\text{mol e}^- \text{ m}^{-2} \text{ s}^{-1}$) | $G_s$ at ambient<br>($\text{mol m}^{-2} \text{ s}^{-1}$) |
| --- | --- | --- | --- | --- |
| Wild type | 5 | $78.2 \pm 6.3$ | $100.10 \pm 5.8$ | $0.17 \pm 0.03$ |
| Line 1 | 7 | $16.9 \pm 2.8$ | $44.5 \pm 7.2$ | $0.18 \pm 0.05$ |
| Line 2 | 7 | $26.1 \pm 4.9$ | $59.8 \pm 11.3$ | $0.25 \pm 0.04$ |
| Line 3 | 7 | $22.7 \pm 4.7$ | $50.8 \pm 11.08$ | $0.16 \pm 0.05$ |

$V_{cmax}$ , maximum rate of Rubisco carboxylation; $J_{max}$ , maximum rate of electron transport; $G_s$ , stomatal conductance at ambient $\text{CO}_2$ . Parameters estimated by fitting $A/C_i$ curves to the Farquhar–von Caemmerer–Berry model (Farquhar et al., 1980) using kinetic constants from Bernacchi et al. (2001). Values are mean ± SD. $G_s$ measured at ambient $\text{CO}_2$ ( $\sim 400 \mu\text{mol mol}^{-1}$ ).

## Notes

### Competing Interest Statement

The authors have declared no competing interest.

## References

Adler L, Díaz-Ramos A, Mao Y, Pukacz KR, Fei C, McCormick AJ (2022) New horizons for building pyrenoid-based CO2-concentrating mechanisms in plants to improve yields. Plant Physiol 190: 1609–1627

Ainsworth EA, Long SP (2021) 30 years of free-air carbon dioxide enrichment (FACE): What have we learned about future crop productivity and its potential for adaptation? Glob Chang Biol 27: 27–49

Atkinson N, Feike D, Mackinder LCM, Meyer MT, Griffiths H, Jonikas MC, Smith AM, McCormick AJ (2016) Introducing an algal carbon-concentrating mechanism into higher plants: location and incorporation of key components. Plant Biotechnol J 14: 1302–1315

Atkinson N, Leitão N, Orr DJ, Meyer MT, Carmo-Silva E, Griffiths H, Smith AM, McCormick AJ (2017) Rubisco small subunits from the unicellular green alga Chlamydomonas complement Rubisco-deficient mutants of Arabidopsis. New Phytologist 214: 655–667

Atkinson N, Mao Y, Chan KX, McCormick AJ (2020) Condensation of Rubisco into a proto-pyrenoid in higher plant chloroplasts. Nat Commun 11: 6303

Atkinson N, Stringer R, Mitchell SR, Seung D, McCormick AJ (2024) SAGA1 and SAGA2 promote starch formation around proto-pyrenoids in Arabidopsis chloroplasts. Proceedings of the National Academy of Sciences 121: e2311013121

Bally J, Jung H, Mortimer C, Naim F, Philips JG, Hellens R, Bombarely A, Goodin MM, Waterhouse PM (2018) The rise and rise of Nicotiana benthamiana: A plant for all reasons. Annu Rev Phytopathol 56: 405–426

Barrett J, Girr P, Mackinder LCM (2021) Pyrenoids: CO2-fixing phase separated liquid organelles. Biochimica et Biophysica Acta (BBA) - Molecular Cell Research 1868: 118949

Bernacchi CJ, Long SP, Ort DR (2025) Safeguarding crop photosynthesis in a rapidly warming world. Science (1979) 388: 1153–1160

Bernacchi CJ, Singsaas EL, Pimentel C, Portis Jr AR, Long SP (2001) Improved temperature response functions for models of Rubisco-limited photosynthesis. Plant Cell Environ 24: 253–259

Bouvier JW, Emms DM, Kelly S (2024) Rubisco is evolving for improved catalytic efficiency and CO2 assimilation in plants. Proceedings of the National Academy of Sciences 121: e2321050121

Clemente T (2006) Nicotiana (Nicotiana tobaccum, Nicotiana benthamiana). *In* K Wang, ed, Agrobacterium Protocols. Humana Press, Totowa, NJ, pp 143–154

Dudley QM, Jo S, Guerrero DAS, Chhetry M, Smedley MA, Harwood WA, Sherden NH, O’Connor SE, Caputi L, Patron NJ (2022) Reconstitution of monoterpene indole alkaloid biosynthesis in genome engineered Nicotiana benthamiana. Commun Biol 5: 949

Duursma RA (2015) Plantecophys - An R Package for Analysing and Modelling Leaf Gas Exchange Data. PLoS One 10: e0143346-

Ellison EE, Nagalakshmi U, Gamo ME, Huang P, Dinesh-Kumar S, Voytas DF (2020) Multiplexed heritable gene editing using RNA viruses and mobile single guide RNAs. Nat Plants 6: 620–624

Engel BD, Schaffer M, Kuhn Cuellar L, Villa E, Plitzko JM, Baumeister W (2015) Native architecture of the Chlamydomonas chloroplast revealed by in situ cryo-electron tomography. Elife 4: e04889

Farquhar GD, von Caemmerer S, Berry JA (1980) A biochemical model of photosynthetic CO2 assimilation in leaves of C3 species. Planta 149: 78–90

Fei C, Wilson AT, Mangan NM, Wingreen NS, Jonikas MC (2022) Modelling the pyrenoid-based CO2-concentrating mechanism provides insights into its operating principles and a roadmap for its engineering into crops. Nat Plants 8: 583–595

Freeman Rosenzweig ES, Xu B, Kuhn Cuellar L, Martinez-Sanchez A, Schaffer M, Strauss M, Cartwright HN, Ronceray P, Plitzko JM, Förster F, et al (2017) The Eukaryotic CO2-Concentrating Organelle Is Liquid-like and Exhibits Dynamic Reorganization. Cell 171: 148–162.e19

Hanson DT, Franklin LA, Samuelsson G, Badger MR (2003) The Chlamydomonas reinhardtii cia3 mutant lacking a thylakoid lumen-localized carbonic anhydrase Is limited by CO2 Supply to Rubisco and not photosystem II function in vivo. Plant Physiol 132: 2267–2275

He S, Chou H-T, Matthies D, Wunder T, Meyer MT, Atkinson N, Martinez-Sanchez A, Jeffrey PD, Port SA, Patena W, et al (2020) The Structural Basis of Rubisco Phase Separation in the Pyrenoid. Nat Plants 6: 1480–1490

He S, Crans VL, Jonikas MC (2023) The pyrenoid: The eukaryotic CO2-concentrating organelle. Plant Cell 35: 3236–3259

He S, Lemma LM, Martinez-Calvo A, He G, Hennacy JH, Wang L, Ergun SL, Rai AK, Wang C, Bunday L, et al (2026) Kinase KEY1 controls pyrenoid condensate size throughout the cell cycle by disrupting phase separation interactions. Nat Cell Biol 28: 725–738

Hennacy JH, Atkinson N, Kayser-Browne A, Ergun SL, Franklin E, Wang L, Eicke S, Kazachkova Y, Kafri M, Fauser F, et al (2024) SAGA1 and MITH1 produce matrix-traversing membranes in the CO2-fixing pyrenoid. Nat Plants. doi: 10.1038/s41477-024-01847-0

Hennacy JH, Jonikas MC (2020) Prospects for engineering biophysical CO2 concentrating mechanisms into land plants to enhance yields. Annu Rev Plant Biol 71: 461–485

Jamet E, Parmentier Y, Durr A, Fleck J (1991) Genes encoding the small subunit of RUBISCO belong to two highly conserved subfamilies in Nicotianeae. J Mol Evol 33: 226–236

Kajala K, Covshoff S, Karki S, Woodfield H, Tolley BJ, Dionora MJA, Mogul RT, Mabilangan AE, Danila FR, Hibberd JM, et al (2011) Strategies for engineering a two-celled C4 photosynthetic pathway into rice. J Exp Bot 62: 3001–3010

Kanevski I, Maliga P (1994) Relocation of the plastid rbcL gene to the nucleus yields functional ribulose-1,5-bisphosphate carboxylase in tobacco chloroplasts. Proceedings of the National Academy of Sciences 91: 1969–1973

Kazachkova Y, Zemach I, Panda S, Bocobza S, Vainer A, Rogachev I, Dong Y, Ben-Dor S, Veres D, Kanstrup C, et al (2021) The GORKY glycoalkaloid transporter is indispensable for preventing tomato bitterness. Nat Plants 7: 468–480

Klein AP, Sattely ES (2017) Biosynthesis of cabbage phytoalexins from indole glucosinolate. Proceedings of the National Academy of Sciences 114: 1910–1915

Lacoste-Royal G, Gibbs SP (1987) Immunocytochemical Localization of Ribulose-1,5-Bisphosphate Carboxylase in the Pyrenoid and Thylakoid Region of the Chloroplast of Chlamydomonas reinhardtii. Plant Physiol 83: 602–606

Laterre R, Pottier M, Remacle C, Boutry M (2017) Photosynthetic Trichomes Contain a Specific Rubisco with a Modified pH-Dependent Activity. Plant Physiol 173: 2110–2120

Lau W, Sattely ES (2015) Six enzymes from mayapple that complete the biosynthetic pathway to the etoposide aglycone. Science (1979) 349: 1224–1228

Leliaert F, Smith DR, Moreau H, Herron MD, Verbruggen H, Delwiche CF, De Clerck O (2012) Phylogeny and Molecular Evolution of the Green Algae. CRC Crit Rev Plant Sci 31: 1–46

Littlejohn GR, Mansfield JC, Christmas JT, Witterick E, Fricker MD, Grant MR, Smirnoff N, Everson RM, Moger J, Love J (2014) An update: improvements in imaging perfluorocarbon-mounted plant leaves with implications for studies of plant pathology, physiology, development and cell biology. Front. Plant Sci. Volume 5–2014:

Long SP, Marshall-Colon A, Zhu X-G (2015) Meeting the Global Food Demand of the Future by Engineering Crop Photosynthesis and Yield Potential. Cell 161: 56–66

Mackinder LCM, Meyer MT, Mettler-Altmann T, Chen VK, Mitchell MC, Caspari O, Freeman Rosenzweig ES, Pallesen L, Reeves G, Itakura A, et al (2016) A repeat protein links Rubisco to form the eukaryotic carbon-concentrating organelle. Proceedings of the National Academy of Sciences 113: 5958–5963

McGrath JM, Long SP (2014) Can the Cyanobacterial Carbon-Concentrating Mechanism Increase Photosynthesis in Crop Species? A Theoretical Analysis. Plant Physiol 164: 2247–2261

Meyer MT, Genkov T, Skepper JN, Jouhet J, Mitchell MC, Spreitzer RJ, Griffiths H (2012) Rubisco small-subunit α-helices control pyrenoid formation in Chlamydomonas. Proceedings of the National Academy of Sciences 109: 19474–19479

Meyer MT, Itakura AK, Patena W, Wang L, He S, Emrich-Mills T, Lau CS, Yates G, Mackinder LCM, Jonikas MC (2020) Assembly of the algal CO2-fixing organelle, the pyrenoid, is guided by a Rubisco-binding motif. Sci Adv 6: eabd2408

Ort DR, Merchant SS, Alric J, Barkan A, Blankenship RE, Bock R, Croce R, Hanson MR, Hibberd JM, Long SP, et al (2015) Redesigning photosynthesis to sustainably meet global food and bioenergy demand. Proceedings of the National Academy of Sciences 112: 8529–8536

Pronina NA, Semenenko VE (1990) Membrane-Bound Carbonic Anhydrase Takes Part in CO2 Concentration in Algae Cells. *In* M Baltscheffsky, ed, Current Research in Photosynthesis: Proceedings of the VIIIth International Conference on Photosynthesis Stockholm, Sweden, August 6–11, 1989. Springer Netherlands, Dordrecht, pp 3283–3286

Raven JA (1997) CO2-concentrating mechanisms: a direct role for thylakoid lumen acidification? Plant Cell Environ 20: 147–154

Raven JA, Cockell CS, De La Rocha CL (2008) The evolution of inorganic carbon concentrating mechanisms in photosynthesis. Philosophical Transactions of the Royal Society B: Biological Sciences 363: 2641–2650

Sage RF, Sage TL, Kocacinar F (2012) Photorespiration and the Evolution of C4 Photosynthesis. Annu Rev Plant Biol 63: 19–47

Sarrion-Perdigones A, Vazquez-Vilar M, Palací J, Castelijns B, Forment J, Ziarsolo P, Blanca J, Granell A, Orzaez D (2013) GoldenBraid 2.0: A comprehensive DNA assembly framework for plant synthetic biology. Plant Physiol 162: 1618–1631

Schindelin J, Arganda-Carreras I, Frise E, Kaynig V, Longair M, Pietzsch T, Preibisch S, Rueden C, Saalfeld S, Schmid B, et al (2012) Fiji: an open-source platform for biological-image analysis. Nat Methods 9: 676–682

da Silva GE, Obst S, Carvalho P, Forner J, Ruf S, Saibo NJM, Bock R (2026) Generation of a recipient line for Rubisco engineering by multiplex genome editing in tobacco. The Plant Journal 126: e70930

Tcherkez GGB, Farquhar GD, Andrews TJ (2006) Despite slow catalysis and confused substrate specificity, all ribulose bisphosphate carboxylases may be nearly perfectly optimized. Proceedings of the National Academy of Sciences 103: 7246–7251

Toyokawa C, Yamano T, Fukuzawa H (2020) Pyrenoid Starch Sheath Is Required for LCIB Localization and the CO2-Concentrating Mechanism in Green Algae. Plant Physiol 182: 1883–1893

Vladimirova MG, Markelova AG, Semenenko VE (1982) Use of the cytoimmunofluorescent method to clarify localization of ribulose bisphosphate carboxylase in pyrenoids of unicellular algae. Soviet Plant Physiology 5: 725–734

Vogan PJ, Sage RF (2011) Water-use efficiency and nitrogen-use efficiency of C3-C4 intermediate species of Flaveria Juss. (Asteraceae). Plant Cell Environ 34: 1415–1430

Wang J, Zhang Q, Tung J, Zhang X, Liu D, Deng Y, Tian Z, Chen H, Wang T, Yin W, et al (2024) High-quality assembled and annotated genomes of Nicotiana tabacum and Nicotiana benthamiana reveal chromosome evolution and changes in defense arsenals. Mol Plant 17: 423–437

Wunder T, Cheng SLH, Lai S-K, Li H-Y, Mueller-Cajar O (2018) The phase separation underlying the pyrenoid-based microalgal Rubisco supercharger. Nat Commun 9: 5076

Zhang Y, Chen M, Siemiatkowska B, Toleco MR, Jing Y, Strotmann V, Zhang J, Stahl Y, Fernie AR (2020) A Highly Efficient Agrobacterium-Mediated Method for Transient Gene Expression and Functional Studies in Multiple Plant Species. Plant Commun 1: 100028

